# HIV-1 *gag-pol* mRNA localization regulates the site of virion assembly

**DOI:** 10.1101/057299

**Authors:** Jordan T. Becker, Nathan M. Sherer

**Author notes:** To whom correspondence should be addressed: 501 Robert M. Bock Lab, 1525 Linden Drive, Madison, WI 53706.

## Abstract

HIV-1 full-length, unspliced genomic RNAs (gRNAs) serve both as mRNAs encoding the Gag and Gag-Pol capsid proteins as well as the genetic material packaged by Gag into virions that assemble at the plasma membrane (PM). Whether localized Gag synthesis contributes to assembly at the PM is unknown. Here we show that artificially tethering gRNAs or surrogate *gag*-pol mRNAs to non-PM membranes or the actin cytoskeleton can markedly affect Gag’s distribution in the cytoplasm, causing aberrant subcellular sites of assembly and severe reductions to virus particle output. Only *gag-pol* mRNAs competent for translation were capable of altering Gag’s distribution within the cell, and the activity mapped to two *cis*-acting RNA regulatory elements; the 5’ packaging signal (Psi) bound by Gag and, unexpectedly, the Rev response element (RRE) that regulates the nuclear export of gRNAs and other intron-retaining viral RNAs. Taken together, our results suggest a model wherein localized translation of gRNAs at the PM helps to compartmentalize Gag-gRNA interactions, thereby promoting efficient genome encapsidation.

**AUTHOR SUMMARY:** The spatial distribution of messenger RNAs (mRNAs) within the cytoplasm can be a crucial determinant of gene expression. Here we provide evidence that a devastating viral pathogen, human immunodeficiency virus type 1 (HIV-1), exploits localized translation to favor the formation of infectious, transmissible virions at the surface of infected cells. Artificially tethering viral mRNAs encoding the Gag and Gag-Pol capsid proteins (*gag-pol* mRNAs) to alternative regions of the cell such as cytoplasmic vesicles or the actin cytoskeletion markedly alters Gag subcellular distribution, perturbs sites of assembly, and reduces virus particle production. These and additional findings suggest a model for HIV-1 assembly wherein localized Gag/Gag-Pol translation coupled to confined interactions between Gag and viral genomes ensures infectious virion production at the right place and the right time. Perturbing HIV-1 mRNA subcellular localization could represent a novel antiviral strategy.

## INTRODUCTION

The spatial distribution of messenger RNAs (mRNAs) within the cytoplasm is a core determinant of mRNA turnover, cytoplasmic utilization, and the formation of functional macromolecular complexes [1–3]. Viruses face severe challenges in this regard during the productive phases of infection wherein viral mRNAs, genomes, and core structural elements must be successfully compartmentalized in space and time to ensure the efficient assembly and release of infectious virions [4–6].

For the retrovirus human immunodeficiency virus type 1 (HIV-1), virion assembly is coordinated at the cytoplasmic face of the plasma membrane (PM) where a dimer of ~9kb, unspliced genomic RNA (gRNA) is encapsidated into an enveloped, proteinaceous shell consisting of ~2,000 Gag (and Gag-Pol) capsid polyproteins [7,8]. To initiate assembly, Gag interacts with the PM via a fatty acid myristoyl membrane anchor and must also interact with an RNA scaffold in the cytoplasm [9–13]. Four functional domains of the 55 kDa Gag precursor polyprotein (Pr55^Gag^) coordinate this process. Matrix (MA/p17^Gag^) targets Gag to the cytosolic face of the PM through interactions with the phospholipid phosphatidylinositol 4,5-bisphosphate (PI(4,5)P_2_) [14–16]. Capsid (CA/p24^Gag^) coordinates Gag-Gag interactions during capsid assembly [17–19,7]. Nucleocapsid (NC/p7^Gag^) binds to gRNAs and/or cellular RNAs [20–24]. The late domain (p6^Gag^) recruits the cellular endosomal sorting complex required for transport (ESCRT) machinery that catalyzes membrane abscission and particle release [25–28].

Upstream of assembly, a single pool of HIV-1 gRNA molecules is thought to serve both as mRNAs encoding Gag and Gag-Pol as well as the core genetic substrate bound by Gag and packaged into virions [29–33]. Gag is translated on free polysomes either prior to or coincident with the formation of viral ribonucleoprotein (vRNP) trafficking granules that consist of low-order multiples of Gag bound to gRNAs in conjunction with cellular RNA binding proteins [34–39]. Recent advanced imaging studies have demonstrated that vRNPs diffuse in the cytoplasm prior to being tethered to the PM by Gag [13,40,41]. Gag-membrane binding is initiated when cytoplasmic Gag concentrations reach a critical level known as the cooperative threshold [42–44], triggering the activation of a myristoyl switch mechanism within MA that subsequently anchors gRNP complexes to the PM [45,46,12].

Gag selectively encapsidates a dimer of gRNA molecules with high efficiency [47–50] due to the NC domain’s capacity to bind a *cis*-acting RNA packaging signal known as *Psi* located in the gRNA’s 5’ untranslated region (UTR) with high affinity [51,21,31,52,53,33,54]. A second cis-acting RNA structure, the Rev response element (RRE), may also contribute to the efficiency of gRNA encapsidation albeit through an unknown mechanism [55,56]. Indeed, the RRE is much better characterized as regulating gRNA nucleocytoplasmic transport through recruitment of the viral Rev protein and subsequent Rev-mediated interactions with cellular CRM1 nuclear export receptor [57–59]. The dimerization of gRNAs may occur in the cytoplasm [60] and/or after Gag-PM anchoring [61], followed by the gradual recruitment of additional Gag molecules to form an immature capsid lattice over a time period of 10-60 minutes [13,62,63,49,41]. NC also binds to and packages cellular RNAs, with highly structured RNAs such as U6 snRNAs and 7SL RNAs encapsidated into virions with a high degree of specificity [64,23,50,65,24]. Interestingly, the MA domain has also been reported to bind RNAs, in particular cellular tRNAs, an activity that regulates MA-membrane interactions *in vitro* and predicted to impact assembly efficiency in cells [53,66,67].

Gag is sufficient to drive the assembly of non-infectious virus-like particles (VLPs) even in the absence of packageable gRNAs [68–73]. Thus, capsid-genome interactions are clearly not obligatory for HIV-1 assembly, unlike for many other viruses [74–76]. On the other hand, imaging studies have demonstrated that gRNAs accumulate at the membrane with Gag prior to the onset of higher-order assembly, so that the gRNA may be capable of nucleating assembly [13,41]. Moreover, we and others have demonstrated that manipulating gRNA trafficking (*e.g.*, rendering gRNAs or surrogate *gag-pol* mRNAs Rev/RRE-independent) can, in some instances, profoundly affect Gag’s capacity to traffic to the PM [77,35,78–81]. An attractive, long-standing hypothesis for links between HIV-1 mRNA trafficking and assembly is that, under native conditions, gRNAs encode one or more signals that influence gRNA/Gag subcellular distribution in the cytoplasm [77,35,80,82]. However, direct evidence for such an activity remains elusive.

In the current study we combined imaging and functional assays to determine if the cytoplasmic abundance or subcellular site of gRNA (*gag-pol* mRNA) localization are determinants of the assembly pathway. Altering gRNA cytoplasmic abundance, in a non-coding context, had little to no stimulatory effect on assembly when provided to Gag in *trans*. By contrast, disrupting gRNA (*gag-pol* mRNA) diffusion in the cytoplasm by artificially tethering gRNAs to non-PM membranes or the actin cytoskeleton markedly affected Gag subcellular distribution and potently reduced virus particle production. Interestingly, these effects were only observed for gRNAs competent for Gag synthesis (*i.e*., *gag-pol* mRNAs), with the effects mapping to the 5’ *Psi* element bound by Gag as well as the Rev response element (RRE) that governs gRNA nuclear export. Taken together, our results are consistent with a model wherein localized Gag translation and compartmentalized Gag-gRNA interactions at the PM promote efficient gRNA encapsidation.

## RESULTS

### Tracking Gag/gRNA interactions in single living cells

To study Gag/gRNA interactions functionally and using fluorescence microscopy, we inserted 24 copies of the MS2 bacteriophage RNA stem-loop (MS2 stem loops; MSL), recognized by the MS2 coat protein, between the *gag* and *pol* open reading frames within the major intron of a full-length HIV-1 NL4-3-based luciferase reporter virus construct (WT-MSL) (Fig 1). WT-MSL expressed full-length Gag and yielded robust production of virus-like particles (VLPs), albeit in the absence of Gag cleavage due to insertion of the MSL cassette upstream of the *pol* gene thereby abolishing synthesis of the viral protease (Figs 1A and 1B, lane 2). In order to visualize gRNAs in living cells, MSL-bearing gRNAs were monitored in HeLa cells engineered to stably express the MS2-YFP protein fused to a carboxy-terminal nuclear localization signal (NLS) (HeLa.MS2-YFP) (Fig 1C). In these cells, low levels of the MS2-YFP protein are sequestered in the nucleus until bound to an MSL-containing gRNA and exported to the cytoplasm. We have previously validated this strategy as a reliable way to obtain direct, single cell measurements of viral mRNA nuclear export, cytoplasmic trafficking behaviors, and translation [82].

**Fig 1.**
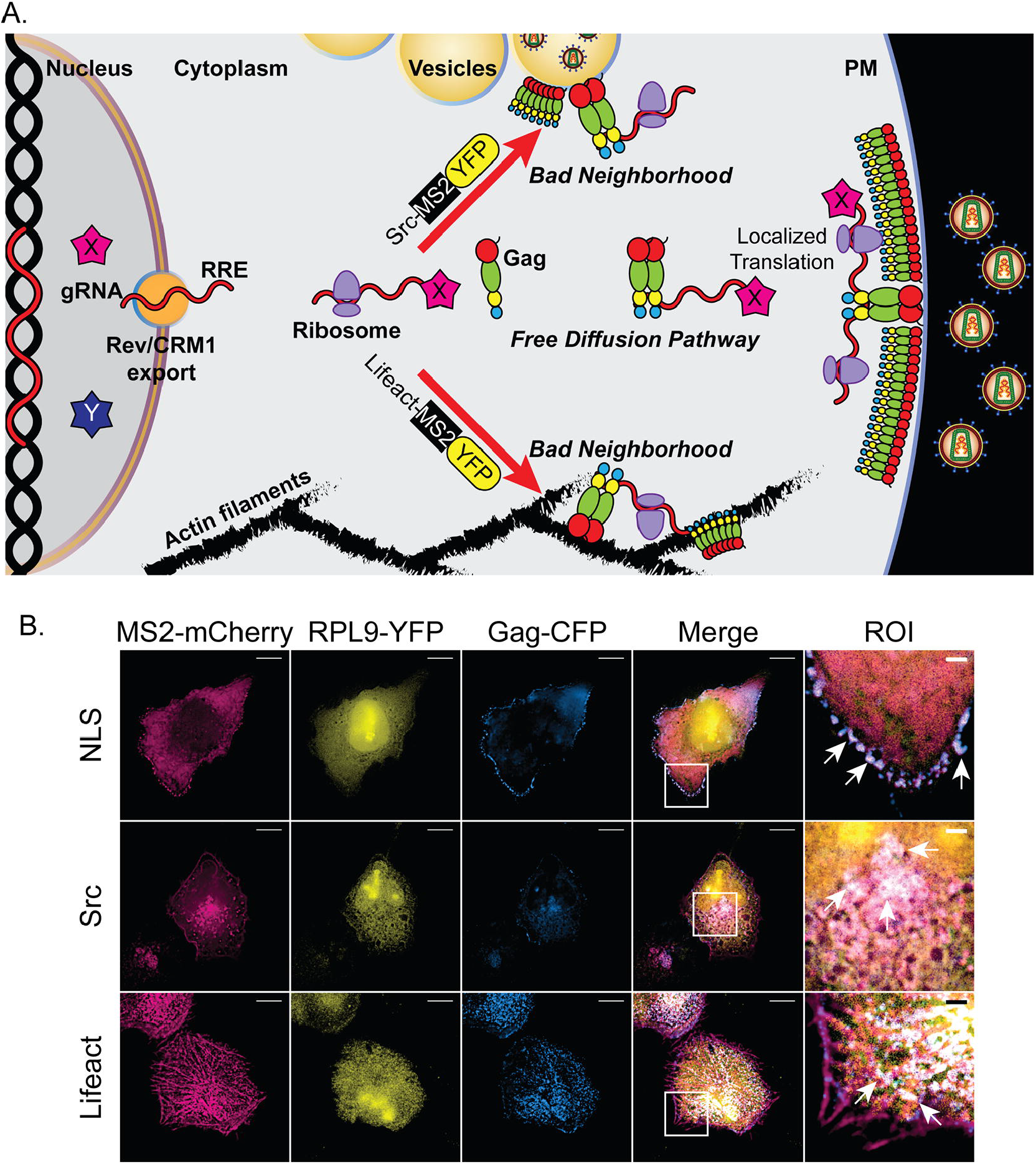
Tracking Gag/gRNA interactions in single living cells. (A) Cartoon depiction of gRNAs used in these studies. Ψ = *Psi* packaging signal. MSL = 24 copies of MS2 RNA stem loop. RRE = Rev-response element. *Tat* and *rev* encode gene-regulatory proteins translated from multiply-spliced mRNAs. *Vif* and *vpu* encode immune modulatory proteins translated from singly-spliced mRNAs. (B) HEK293T cells were transfected with 2000ng HIV-1 plasmids encoding the WT, modified, and mutated gRNAs depicted in (A). VLPs and cell lysates were collected 48 hours post-transfection, resolved by SDS-PAGE, and detected by immunoblot using anti-p24^Gag^ antiserum (HSP90 was also detected as a loading control). (C) Widefield deconvolution microscopy images of HeLa.MS2-YFP cells transfected with 100ng RevInd GagFP and 900ng HIV constructs, fixed ~30 hours post-transfection, and imaged. Single Z-plane images are shown. Scale bars represent 10 microns in full images, 2 microns in regions of interest (ROI). Dashed white lines show the relative position of cell nuclei. White box outlines the ROI. Red arrows indicate sites where RevInd GagFP has accumulated in PM-adjacent punctae. (D) Quantification of MS2-YFP localization phenotypes. Bar graphs show percent of transfected cells with nuclear, cytoplasmic, or both distributions of gRNA for each transfection condition. (E) Quantification of GagFP distribution phenotypes. Bar graphs show percent of transfected cells with diffuse or PM-adjacent punctae Gag localization for each transfection condition. For both (D) and (E), error bars represent standard deviation from the mean for at least four independent experiments quantifying at least 100 cells per condition. (F) HEK293T cells were co-transfected with 500ng RevInd GagFP and 1500ng of HIV gRNA constructs as indicated or an empty vector control (pBluescript) and immunoblotted as in 1A. Bar graphs show fold change in Gag release factor relative to empty vector condition. Release factor is calculated by Gag band intensities in VLPs divided by lysates normalized to HSP90 (N=4).

Investigating the role of gRNAs during the process of assembly is confounded by the gRNA’s essential role as the *gag/gag-pol* mRNA [35]. Thus, we also monitored “gRNA-only” transcripts bearing a single nucleotide substitution (ATG>ACG) at the initiator methionine codon of Gag (1ACG-MSL, depicted in Fig 1A). To control for cytoplasmic activities, we mutated 1ACG-MSL transcripts by deleting the Rev response element (RRE), thus generating a gRNA incapable of exiting the nucleus due to the inability of HIV-1 Rev to regulate CRM1-dependent gRNA nucleocytoplasmic trafficking (dRRE-MSL, Fig 1A). As expected, neither 1ACG nor dRRE transcripts were competent for the translation of full-length Gag proteins (Fig 1B, compare lanes 3 and 4 to lane 2). However, 1ACG transcripts were both exported from the nucleus (Fig 1C) and translated, as evidenced by the synthesis of low levels of a minor Gag isoform (p40) previously shown to result from initiation at *gag* codon methionine-142 (Fig 1B, lane 3) [83,84]. To supply “gRNA-minus” Gag in *trans*, we expressed Gag-fluorescent protein (FP = cyan fluorescent protein, mTagBFP2, mCherry, *etc.*, depending on the experiment but shown in blue throughout for consistency) fusion proteins from constructs wherein the *gag* coding region was codon-optimized to achieve protein synthesis in the absence of Rev or any other viral factors (RevInd GagFP) [85,86] (Figs 1A and 1B, lane 5).

We first tested if RevInd GagFP trafficking or assembly efficiency was affected by the provision of cytoplasmic HIV-1 gRNAs in *trans*. As expected, expression of WT-MSL and 1ACG-MSL gRNAs yielded translocation of MS2-YFP from the nucleus to the cytoplasm in >50% of transfected HeLa.MS2-YFP cells at ~24 hours post-transfection (Figs 1C and quantification in 1D). By contrast, dRRE-MSL gRNAs formed discrete MS2-YFP punctae that were retained in the nucleus in >90% of transfected cells, consistent with gRNA transcription events but in the absence of nuclear export (Figs 1C and 1D). When co-expressed, WT- or 1ACG-gRNAs co-localized with RevInd GagFP aggregates at the plasma membrane, suggesting Gag/gRNA cotrafficking to assembly sites (Fig 1C, bottom panels). However, for each of these conditions, we observed only minor differences in the frequency of cells exhibiting PM-adjacent RevInd GagFP aggregates (Fig 1C, middle panels, and quantification in 1E). Moreover, RevInd GagFP was released from cells with similar efficiency when co-expressed with either 1ACG- or dRRE-gRNAs in a VLP assembly assay using HEK293T cells (Fig 1F). These experiments demonstrated that RevInd GagFP trafficking is largely unaffected by gRNAs co-expressed in the cytoplasm and accessed in *trans*.

### Increasing HIV-1 gRNA cytoplasmic abundance has only minor effects on GagFP trafficking at low, sub-cooperative levels

Because per cell Gag expression levels vary during transient transfection, we next tested the hypothesis that gRNA cytoplasmic abundance is more relevant to the assembly pathway at low, sub-cooperative levels of Gag using HeLa cells engineered to stably express only low levels of RevInd Gag-CFP (HeLa.Gag-CFP cells). Consistent with previous reports defining cooperative assembly [42,43], these cells did not exhibit marked Gag-CFP fluorescence along the PM or diffraction-limited Gag-CFP punctae at the PM. We selected a high performance clone wherein RevInd Gag-CFP was typically observed in a diffuse distribution throughout the cytoplasm until the cells were infected with an HIV-1/mCherry reporter virus, wherein the Gag-CFP signal was markedly relocalized to punctae at the PM at 24-36 hours post-infection (Fig 2A and S1 Video). This visual detection of the onset of particle production demonstrated the utility of this cell line as a biosensor for detecting HIV-1 assembly activity in real time.

**Fig 2.**
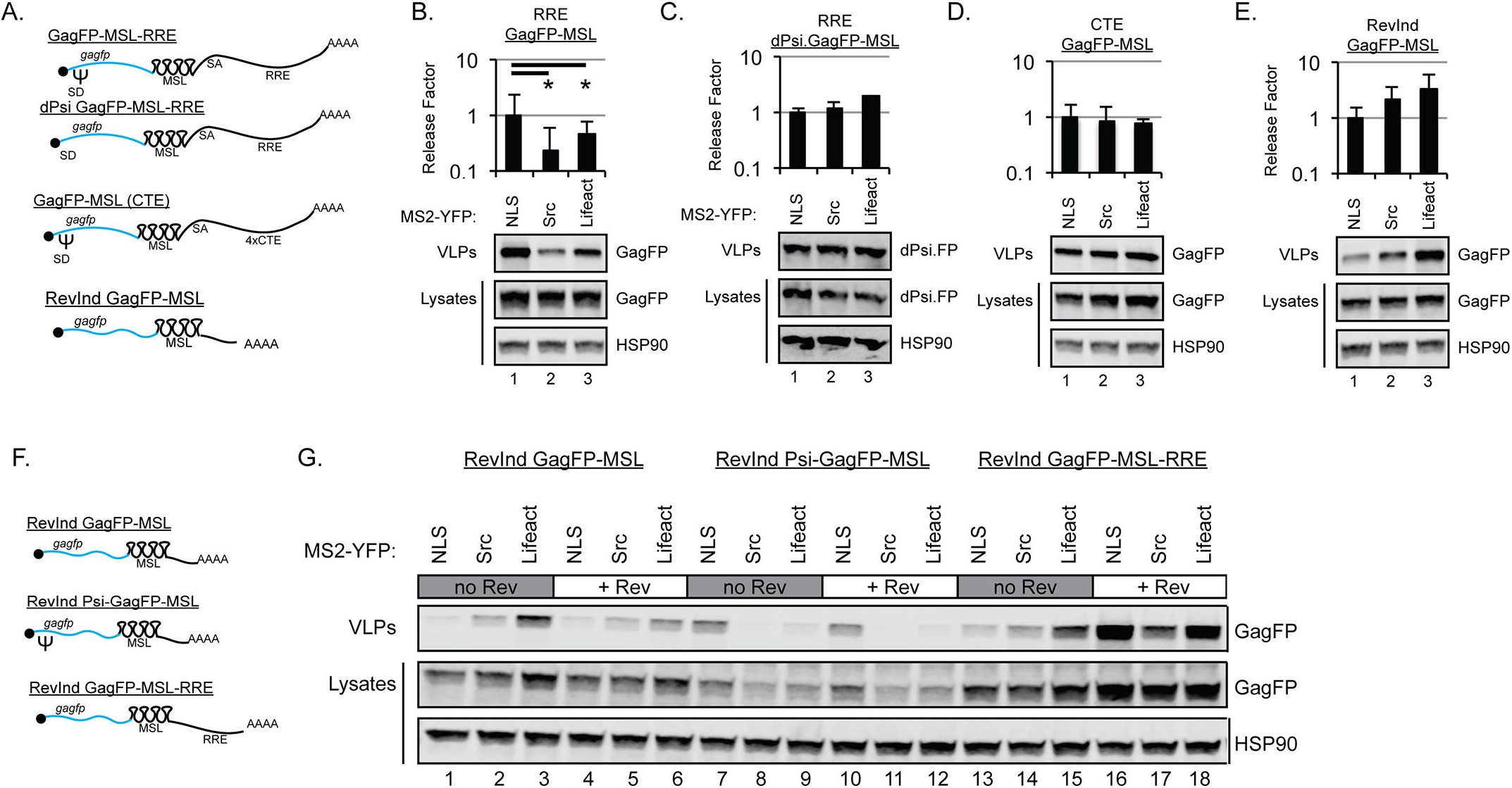
HIV-1 gRNA cytoplasmic abundance plays a minor role in GagFP trafficking at the sub-cooperative threshold. (A) Widefield deconvolution microscopy images from live cell imaging experiments of HeLa.Gag-CFP cells infected with WT NL4-3 E-R-mCherry reporter virus. Multi-channel images were acquired once per hour for up to 48 hours beginning at ~1 hour post-infection. Scale bars represent 10 microns. Red arrows indicate sites where stably expressed RevInd Gag-CFP has transitioned from a diffuse cytoplasmic distribution to PM-associated punctae. (B) Depiction of “self-labeling” WT-MSL gRNA. These gRNAs are identical to those depicted in Fig 1A with the addition of MS2-mCherry-NLS as a reporter and gRNA-tagging protein expressed from the viral *nef* gene position. (C) Widefield deconvolution microscopy images of stable HeLa.Gag-CFP cells transfected with 1000ng HIV gRNA constructs and fixed ~30 hours post-transfection. Scale bars represent 10 µm in full images, 2 µm in ROI. Dashed white lines show the relative position of cell nuclei. The white box designates the ROI. Red arrows indicate sites where stably expressed Gag-CFP has accumulated at the PM for WT-MSL or in the cytoplasm for 1ACG-MSL. dRRE-MSL gRNAs are not exported from the nucleus. Dashed red line in ROI represents edge of cell. (D & E) Quantification of Gag-CFP distribution phenotypes from live cell imaging experiments performed similar to Fig 2A. HeLa.Gag-CFP cells were transfected with 333ng of gRNA constructs (WT-MSL, 1ACG-MSL, and dRRE-MSL also encoding MS2-mCherry-NLS as a reporter and gRNA tagging protein). (D) Bar graphs show the percentage of transfected cells with diffuse or PM-adjacent punctae Gag localization. (E) Bar graph shows the percentage of transfected cells with diffuse, granules, or PM-adjacent punctae Gag localization. Error bars represent standard deviation from the mean. At least 30 cells were quantified per transfection condition per experiment (N=3). (F) HEK293T.Gag-CFP cells were transfected with 2000ng of HIV gRNA constructs as indicated or an empty vector control (pBluescript) and immunoblotted for Gag and HSP90. Bar graphs show fold change in Gag release factor relative to empty vector condition (N=3). The asterisk (*) indicates stable GagFP release factor for WT-MSL condition is significantly different from empty vector condition (Two-tailed Student’s t-test, p=0.0006).

Because the HeLa.Gag-CFP cell line lacked MS2-YFP, in these experiments we tracked Gag-CFP in the presence of WT-MSL, 1ACG-MSL, and dRRE-MSL constructs modified to express MS2-mCherry-NLS from the native viral promoter (*i.e.*, with MS2-mCherry-NLS inserted into the *nef* reading frame, Fig 2B). In HeLa.Gag-CFP cells expressing WT-MSL/MS2-mCherry-NLS gRNAs, we observed marked transitions of RevInd Gag-CFP to discrete punctae at the PM of >75% of cells, identical to the transitions observed after infection (Fig 2A) and thus demonstrating recruitment of Gag-CFP into budding virions (Figs 2C, left panels and quantification in 2D). The gRNA’s MS2-mCherry-NLS signal (yellow) co-localized with surface Gag-CFP (cyan), consistent with Gag/gRNA co-trafficking to the PM and suggesting gRNA encapsidation (Fig 2C, lower left panel). Gag-CFP was also released from cells as VLPs under this condition (Fig 2F, compare lane 2 to lane 1). Contrary to our hypothesis, accumulation of 1ACG-MSL gRNAs in the cytoplasm did not drive RevInd Gag-CFP at sub-cooperative levels to the PM. Instead, we observed RevInd Gag-CFP as well as cytoplasmic MS2-mCherry-NLS signals coalescing in large cytoplasmic granules in >50% of cells (Figs 2C, middle panels, quantification in 2E, and S2 Video). dRRE-MSL expression also had no effect on RevInd Gag-CFP distribution, as expected (Figs 2C, right panels, 2D, and 2E). Taken together, these data indicated that a sub-cooperative threshold for Rev-independent Gag assembly cannot be lowered (or reached) by an excess abundance of cytoplasmic gRNA in single cells, at least when provided in *trans*. In fact, the tendency of 1ACG-gRNAs to aggregate with Gag-CFP in large cytoplasmic granules suggests that disproportionately high levels of gRNA in the cytoplasm are detrimental to Gag trafficking.

### Perturbing the subcellular localization of HIV-1 gRNAs competent for Gag synthesis disrupts virus particle production

The above experiments indicated that altering gRNA cytoplasmic abundance has little to no effect on Gag subcellular trafficking or in nucleating assembly events at the PM when provided in *trans*. We next tested if gRNA subcellular localization is a determinant of the assembly pathway. To this end, we modified MS2-YFP fusion proteins to carry subcellular trafficking motifs in order to artificially target MSL-bearing gRNAs to specific cellular membranes or the actin cytoskeleton (Figs 3–6). We first tested a protein myristoylation signal (MGSSKSKPKD) derived from the proto-oncogene Src kinase, and generated versions of Src-MS2-YFP that would or would not accumulate preferentially in association with the nucleus due to the presence or absence of a carboxy-terminal NLS (Src-MS2-YFP and Src-MS2-YFP-NLS, respectively) (Fig 3A). The Src targeting motif was chosen for these experiments because, similar to Gag’s MA domain, it targets proteins to PI(4,5)P_2_ phospholipid moieties at the cytoplasmic face of the PM [87]. Indeed, prior work has shown that the assembly of Gag mutants lacking MA is rescued by the addition of an amino-terminal Src membrane targeting motif [42,88]. As a control, we employed a previously validated MS2-NXF1 fusion protein that alters gRNA nucleocytoplasmic transport by biasing it toward the NXF1/NXT1 nuclear export pathway, in competition with Rev and CRM1 [89,90]. Fluorescence microscopy confirmed that the Src-MS2-YFP protein localized predominantly to the PM, as expected (Fig 3B). Interestingly, the addition of the NLS (Src-MS2-YFP-NLS) resulted in preferential targeting to the nuclear membrane (Fig 3B), perhaps accessing nucleus-associated PI(4,5)P2 [91].

**Fig 3.**
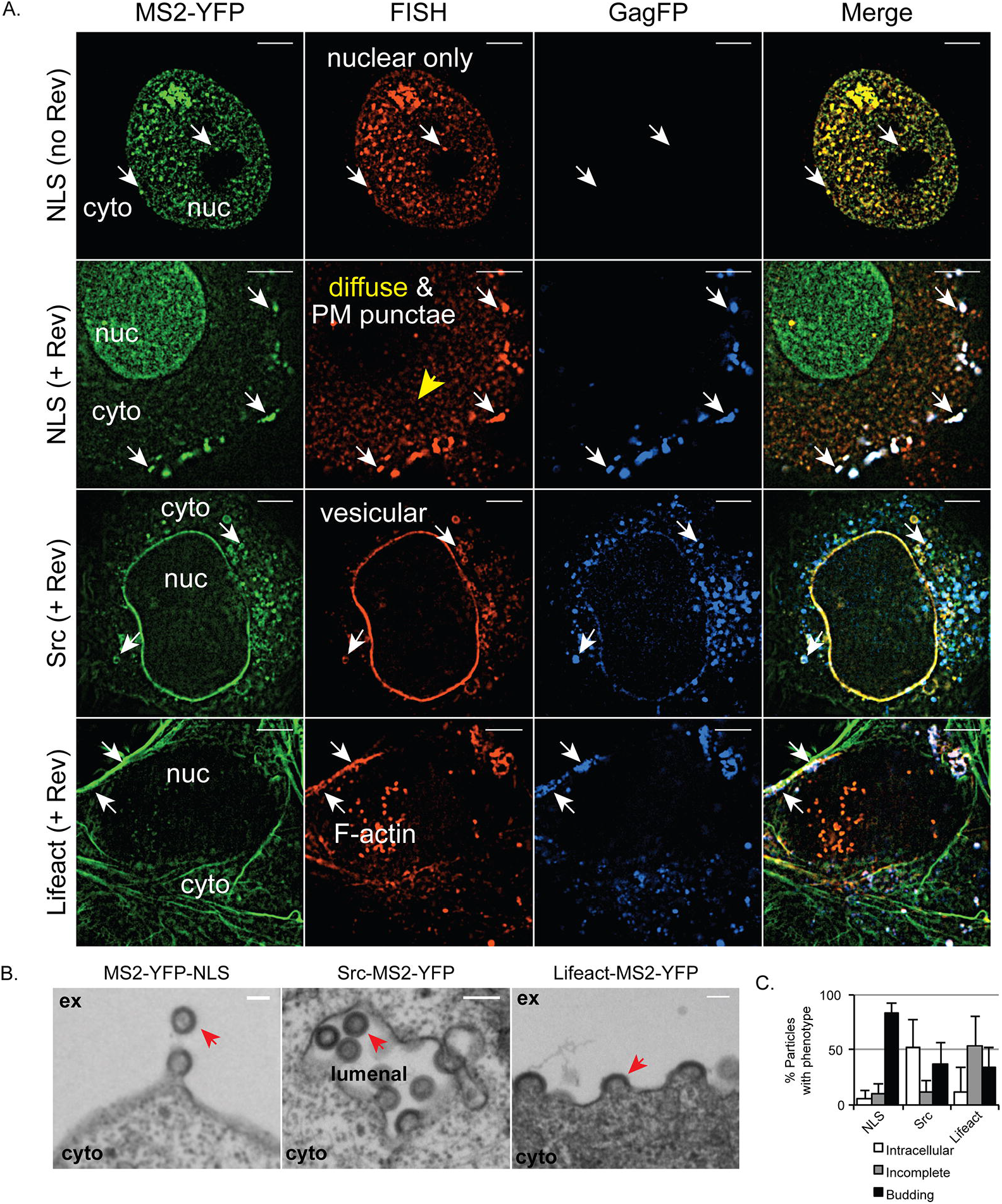
Perturbing HIV-1 gRNA subcellular localization disrupts virus particle production. (A) Cartoon depiction of MS2-YFP targeting protein constructs used in these studies. Short name used in subsequent Figs is underlined. Amino acid targeting motif is shown at their relative (amino- or carboxy-terminal) position. (B) Widefield deconvolution microscopy images of HeLa cells transfected with 333ng MS2-YFP targeting constructs and fixed ~30 hours post-transfection. Scale bars represent 10 µm. Dashed white lines show the relative position of cell nuclei. Dashed cyan lines show the edge of cell. (C) HEK293T cells were transfected with 1000ng MS2-YFP targeting constructs as indicated and 1000ng of WT-MSL and immunoblotted for Gag and HSP90. Bar graphs show release factor relative to Free MS2-YFP. Error bars represent standard deviation from the mean for three independent experiments. The asterisks (*) indicate Pr55 Gag release factor is significantly different for comparisons indicated by black bars (two-tailed Student’s t-test, p=0.023 Src and 0.015 Src+NLS). (D) HEK293T cells were transfected with 1000ng MS2-YFP targeting constructs as indicated and 1000ng of WT NL4-3 E-R- and immunoblotted for Gag and HSP90. Cells were treated with the HIV-1 protease inhibitor saquinavir to prevent Pr55 Gag proteolytic processing and aid quantification of Gag expression and release. Bar graphs show release factor relative to Free MS2-YFP. Error bars represent standard deviation from the mean of three independent experiments. No conditions were significantly different. (E) HEK293T cells were transfected with the indicated amounts of WT NL4-3 E-R- plasmid as indicated and processed as for (D). Bar gr103 aphs show release factor relative to 1000ng condition. Error bars represent standard deviation from the mean for three independent experiments. (C-E). Numbers above blot images represent Gag intensity value of band directly below, relative to lane 1 of each image.

We initially hypothesized that the MS2-YFP protein bearing the Src membrane-targeting signal would stimulate virus particle assembly by enhancing MSL-dependent gRNA trafficking to the PM, and thus provide a nucleation signal to Gag. However, we observed the opposite outcome, with co-expression of either Src-MS2-YFP or Src-MS2-YFP-NLS causing a greater than ten-fold reduction in VLP release for Gag derived from MSL-bearing gRNA (WT-MSL) transcripts (Fig 3C, compare lanes 3 and 4 to lanes 1 and 2). Parental WT gRNAs lacking the MSL cassette were relatively immune to the MS2 targeting proteins (Fig 3D), even at a high ratio (1:1) of MS2:gRNA plasmids, thus demonstrating that the bulk of the effect was specific to MS2-MSL interactions. The MS2-NXF1 control also affected VLP release specifically from the WT-MSL construct. However, this effect was associated with a marked increase to levels of cell-associated Gag, likely reflecting enhanced gRNA nuclear export and/or effects on gRNA trafficking or Gag synthesis intrinsic to the NXF1/NXT1 pathway (Fig 3C, lane 5).

Because assembly is a cooperative process (*i.e.*, highly sensitive to intracellular Gag levels) [12,42,92], we tested if the effects on virus particle production reflected either 1) net reductions to per cell Gag expression levels, 2) decreases to Gag assembly efficiency, or 3) a combination of both effects. In our experiments, we measured virus assembly competency by calculating a “release factor” (RF) [80], defined as the amount of Gag detected in VLPs released into the culture media (determined by quantitative infrared immunoblot), and compared to relative levels of cell-associated Gag. Careful control titrations of the WT construct from 1000 ng to 31.25 ng per transfection (Fig 1E) demonstrated a remarkably linear (R^2^=0.97) relationship between Gag cytoplasmic abundance and VLP production at all levels of Gag in our assays (*i.e.*, a 10-fold loss to VLP production correlated to a 10-fold decrease to cytoplasmic Gag abundance, compare Fig 3E lane 1 to lane 3). For the Src-MS2-YFP conditions, we observed a ~2-fold reduction to cell-associated Gag for both the WT-MSL and WT constructs (e.g., compare Fig 3C lanes 3 and 4 to Fig 3D lane 3). However, the effects on VLP release for the MSL-bearing construct was much greater (>10-fold compared to 3-fold, compare Figs 3C VLPs lanes 3 and 4 to Fig 3D lane 3). Thus, Src-MS2-YFP interactions with WT-MSL gRNAs were apparently affecting not only Gag synthesis but also virus particle release.

### Tethering *gag-pol* mRNAs to non-PM membranes disrupts Gag’s trafficking to the plasma membrane

To address the mechanism underpinning the Src-MS2-YFP effects on Gag abundance and trafficking, we directly monitored Gag in single cells using previously validated, intron-retaining and Rev-dependent GagFP-MSL-RRE surrogate gRNA transcripts [82] (Figs 3–7). Similar to WT-MSL gRNAs, GagFP-MSL-RRE VLP release was completely inhibited by Src-MS2-YFP or Src-MS2-YFP-NLS proteins (Fig 4B, compare lanes 3 and 4 to 1 and 2). Increasing the ratio of Src-MS2-YFP:gRNA plasmids lowered the amount of GagFP released from cells in the form of VLPs without changing levels of cell-associated GagFP-RRE-MSL (Fig 4C, lanes 4–6), a result, again, consistent with a block to virus particle assembly or release from the cell.

**Fig 4.**
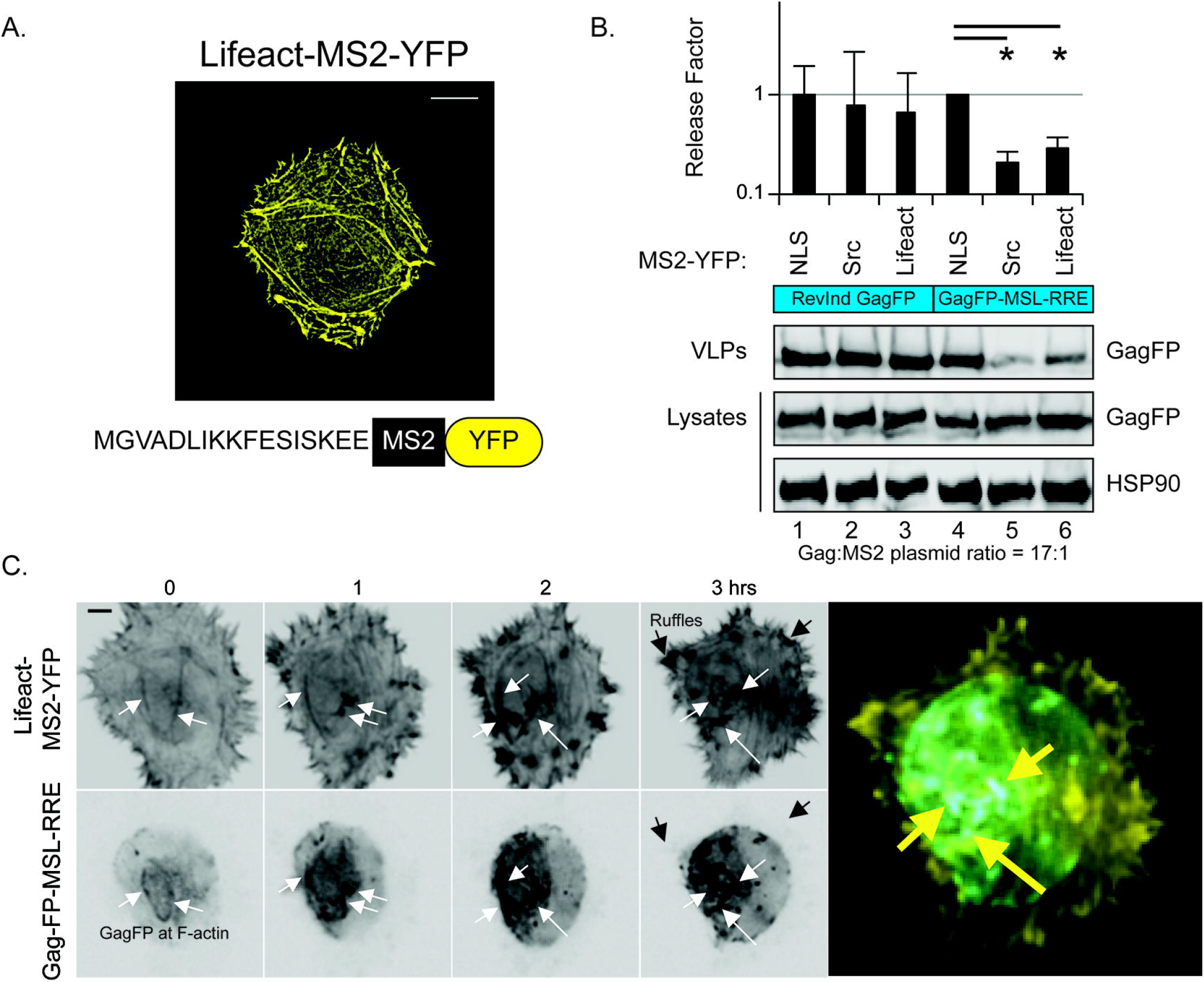
Src-MS2-YFP proteins induce a gRNA-specific block to Gag trafficking in *cis*. (A) Cartoon depiction of subgenomic HIV-1 GagFP-MSL-RRE construct used. Splice donor (SD) and splice acceptor (SA) are shown to emphasize that the viral *gagfp* mRNA (surrogate subgenomic gRNA) retains an intron. (B) HEK293T cells were transfected with 1000ng MS2-YFP targeting constructs as indicated, 900ng of GagFP-MSL-RRE, and 100ng pRev and immunoblotted for Gag and HSP90. Bar graphs show release factor relative to Free MS2-YFP. Error bars represent standard deviation from the mean for three independent experiments. The asterisks (*) indicate Gag release factor is significantly different for comparisons indicated by black bars (two-tailed Student’s t-test, p=0.03). (C) HEK293T were transfected with decreasing amounts of GagFP-MSL-RRE (1500/1000/500ng) plus 200ng Rev and empty vector as filler up to 2µg total DNA in lanes 1-3. Cells were transfected with 1500ng GagFP-MSL-RRE, plus 200ng Rev, empty vector as filler, and increasing amounts (100/200/300ng) of Src-MS2-YFP and immunoblotted for Gag and HSP90 in lanes 4-6. Bar graphs show release factor relative to lane 1. Error bars represent standard deviation from the mean of three independent experiments. (D) *Cis* versus *trans* effects. HEK293T cells were transfected with 500ng RevInd Gag-CFP, 100ng MS2-YFP targeting construct as indicated, and 1400ng empty vector, 1ACG (no MSL), or 1ACG-MSL in lanes 1-6. Lanes 7-8 were transfected with 1400ng GagFP-MSL-RRE, 100ng MS2-YFP targeting construct as indicated, 200ng Rev, and 300ng empty vector immunoblotted for Gag and HSP90. (E) Widefield deconvolution microscopy images of HeLa cells transfected with 100ng MS2-YFP targeting constructs, 800ng subgenomic GagFP-MSL-RRE, and 100ng pRev and fixed ~30 hours post-transfection. Scale bars represent 10 µm in full images and 2 µm in regions of interest (ROI). Dashed white lines show the relative position of cell nuclei. White boxes outline the ROIs. Red arrows indicate sites where GagFP has accumulated. (F) Quantification of GagFP localization phenotypes. Bar graphs show the percentage of cells exhibiting vesicular, diffuse cytoplasmic, or PM-associated puncate for each transfection condition. Error bars represent the standard deviation from the mean for three independent experiments, quantifying at least 100 cells per condition. (G) Whole field quantification of GagFP fluorescence for ~500 cells comparing the permissive NLS and non-permissive Src conditions in cells fixed and imaged at 24 hours post-transfection. HeLa cells were transfected with 150ng MS2-YFP targeting construct, 750ng GagFP-MSL-RRE, and 100ng pRev. Error bars represent the standard deviation from the mean for three independent transfections.

We also tested if Src-MS2-YFP proteins could inhibit VLP production in *trans* by co-transfecting RevInd GagFP with 1ACG genomes either lacking or bearing the MSL cassette, and also in the presence or absence of either control MS2-YFP-NLS or inhibitory Src-MS2-YFP proteins. Src-MS2-YFP did not inhibit assembly by RevInd GagFP under these conditions (Fig 4D, compare lanes 3 and 6 to the controls in lanes 7 and 8), thus indicating that the Src-MS2-YFP-induced, gRNA-dependent assembly inhibition only operates in *cis*, in the context of the *gag/gag-pol* mRNA.

Fluorescence microscopy revealed GagFP to be less frequently detected at PM punctae under these conditions (measured at 24 hours post-transfection), and most often found associated with cytoplasmic vesicles (Fig 4E and quantification in 4F), perhaps consistent with the capacity of the Src-derived trafficking signals to track PI(4,5)P2 throughout the endocytic pathway [91,93]. Fluorescence-based measurements of GagFP demonstrated no differences to cell-associated Gag at this low yet inhibitory ratio of Src-MS2-YFP:gRNA plasmids (1:5) (Figs 4B and fluorescence measured in 4G). Time lapse imaging of both MS2-YFP and GagFP constructs simultaneously in single cells over a 16 hour time course revealed that, for the control MS2-YFP-NLS protein, Gag filled the cytoplasm gradually prior to formation of bright Gag- and gRNA-positive punctae at the PM. This was consistent with the expected punctuated burst of gRNA nuclear export, Gag translation, Gag/gRNA diffusion in the cytoplasm, and ultimately the formation of higher order assembly intermediates at the PM (Fig 5A, black arrows). By contrast, the Src-MS2-YFP protein drove GagFP to accumulate at perinuclear structures, visible even at the lowest levels of GagFP detected (Fig 5B, black arrows). We concluded from these experiments that a convergence of Src-MS2-YFP, gRNA, and Gag leads to the aggregation of Gag/gRNA transport complexes at non-PM membranes, with a possible explanation being Gag accumulating in close proximity to its mRNA and localized site of translation. Such a behavior would not be expected for Gag and gRNA when expressed in *trans* (as suggested by Fig 2 and consistent with Fig 4D).

**Fig 5.**
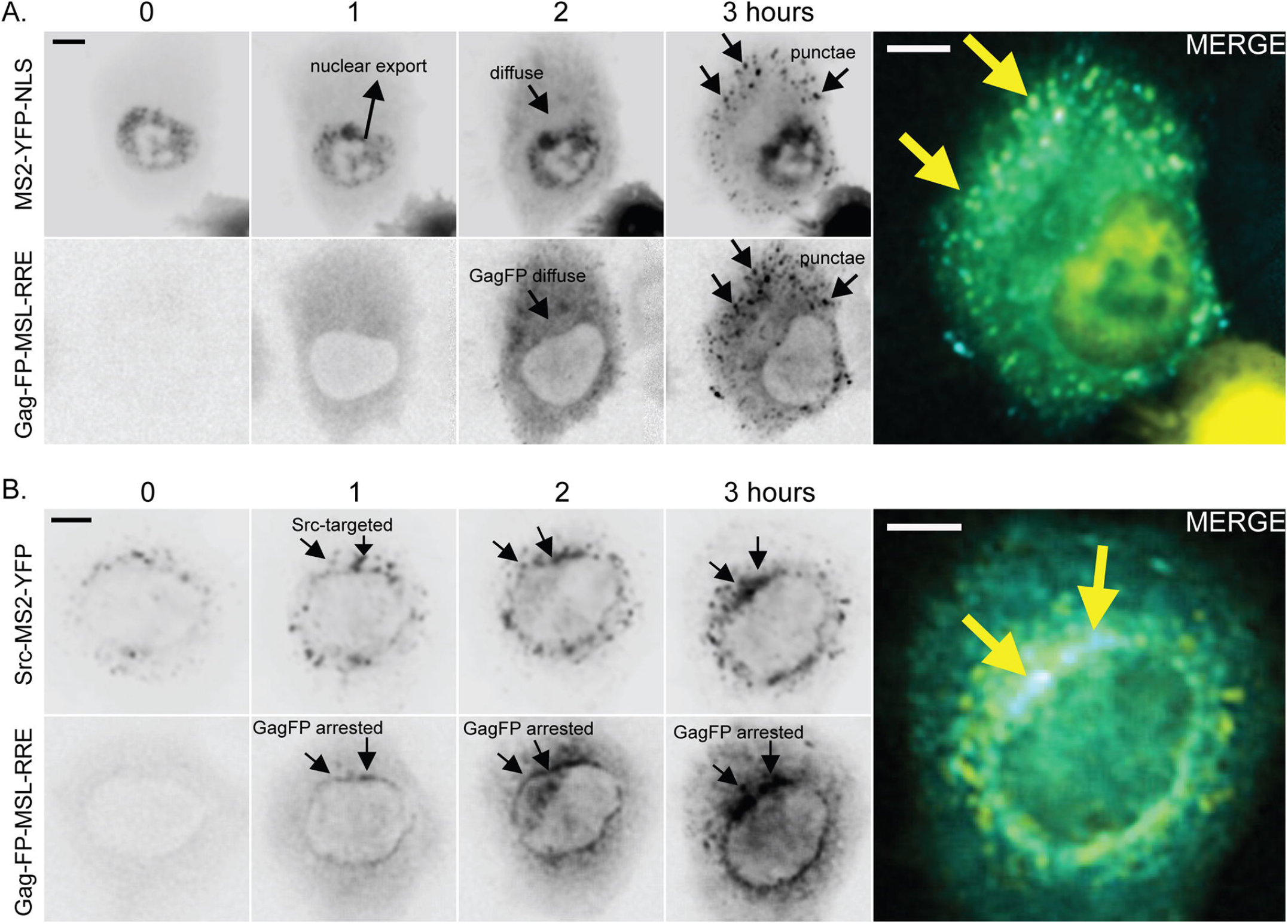
Mis-targeted gRNAs are capable of physically re-routing Gag trafficking to aberrant subcellular locations even at very low levels of Gag expression. (A) Single HeLa cells expressing GagFP derived from GagFP-MSL-RRE constructs, Rev, and MS2-YFP-NLS transfected as for 4E and monitored using live cell fluorescence microscopy for 16 hours. As expected [82], both GagFP and MS2-YFP signals accumulated in the cytoplasm over time prior to coalescing in bright puncate at the PM. (B) When co-expressed with the Src-MS2-YFP protein, both Src-MS2-YFP and GagFP aggregated in a perinuclear zone at even the lowest levels of GagFP detected (see T=0 and T=1h).

### Effects of targeting *gag* mRNAs to the actin cytoskeleton

The experiments above indicated that tethered *gag-pol* mRNAs (as represented by the surrogate subgenomic GagFP gRNA transcripts) physically reposition sites of Gag synthesis from typical cytoplasmic diffusion paths toward the PM (see Fig 5A) to alternative locations (Fig 5B). To further test the capacity of gRNAs to control Gag trafficking, we tested a second targeting protein, Lifeact-MS2-YFP, intended to bias gRNA trafficking to the cell periphery due to its strong interactions with the cortical actin cytoskeleton (Fig 6). Several studies have implicated cortical actin in HIV-1 trafficking/assembly [94–99]. Lifeact is a 17 amino acid peptide capable of targeting proteins to F-actin bundles with high specificity [100] and, as expected, Lifeact-MS2-YFP localized to peripheral actin fibers throughout the cell (Fig 6A). Both Src-MS2-YFP and Lifeact-MS2-YFP had inhibitory effects on assembly (Fig 6B), although Lifeact-MS2-YFP was typically less potent than Src-MS2-YFP (Fig 6B, compare lanes 5 and 6 to 4, and see Fig 8). Two-color time-lapse single cell imaging confirmed that GagFP was rapidly targeted to actin fibers with LifeAct MS2-YFP, again at the earliest and lowest detectable levels of GagFP expression (Fig 6C, white arrows). Interestingly, at early time points GagFP clustered preferentially with Lifeact-MS2-YFP at F-actin bundles at or near the cell body (the rounded portion of the cell) and was less frequently observed in association with dynamic actin ruffles or filopodia at the cell periphery (Fig 5C, black arrows at T=3h). These experiments confirmed that re-targeted *gag-pol* mRNAs are sufficient to target Gag to even relatively exotic subcellular locales such as the actin cytoskeleton.

**Fig 6.**
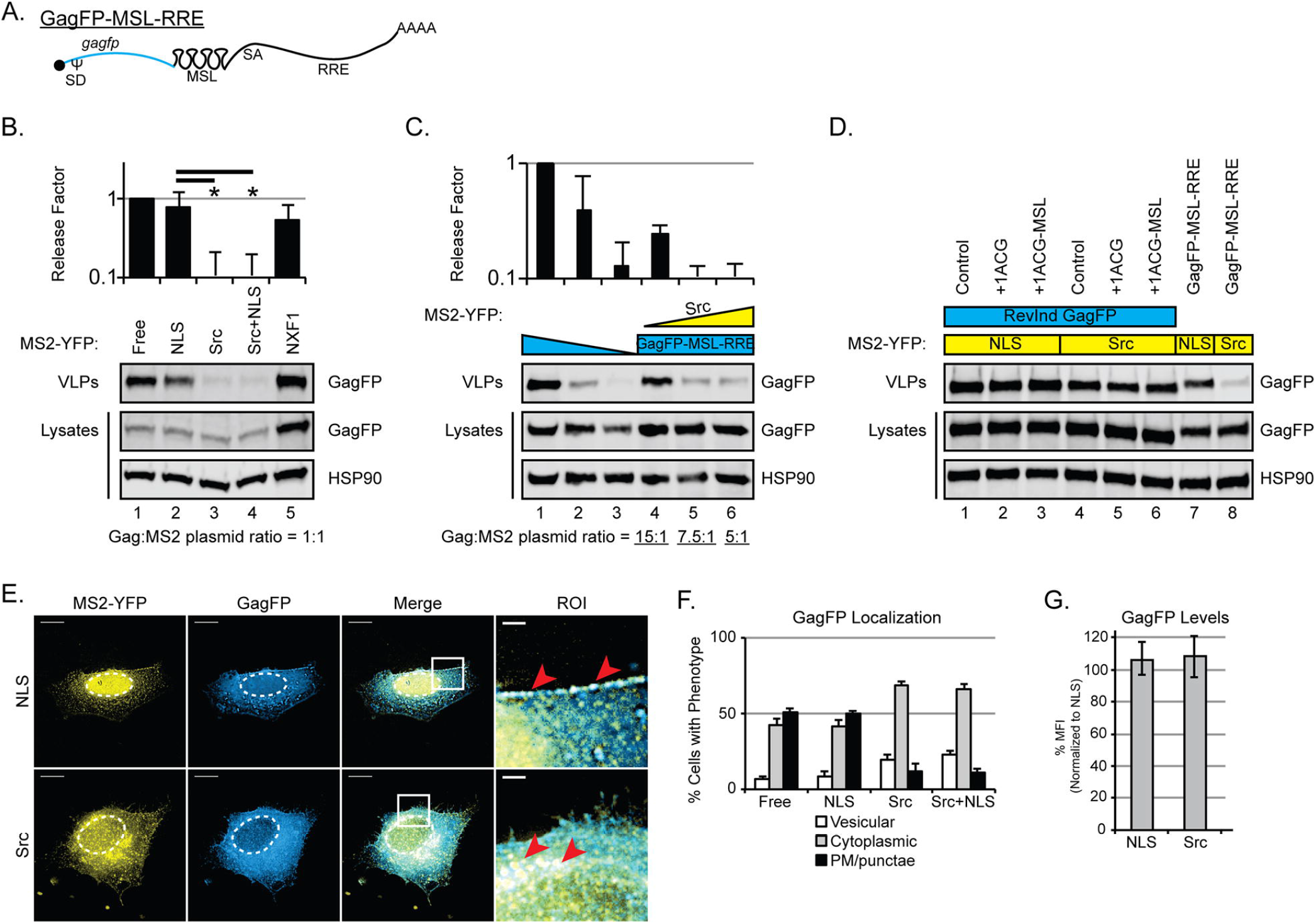
Effects of re-targeting gRNAs to the actin cytoskeleton. (A) Cartoon depiction of Lifeact-MS2-YFP targeting protein construct and widefield deconvolution microscopy image of HeLa cell transfected with 333ng MS2-YFP targeting constructs, fixed ~30 hours post-transfection. Scale bar represent 10 microns. (B) HEK293T cells were transfected with 100ng MS2-YFP targeting constructs as indicated and either 1700ng RevInd GagFP (no MSL) with empty vector OR 1700ng of GagFP-MSL-RRE with 200ng pRev and immunoblotted for Gag and HSP90. Bar graphs show release factor relative to MS2-YFP-NLS condition for each Gag type (RevInd GagFP or GagFP-MSL-RRE). Error bars represent standard deviation from the mean of three independent experiments. The asterisks (*) indicate GagFP release factor is significantly different for comparisons indicated by black bars (two-tailed Student’s t-test, p=0.0001). (C) Images from live cell time-lapse fluorescence microscopy of a HeLa cell expressing GagFP derived from 750ng GagFP-MSL-RRE constructs co-transfected with 100ng Rev and 150ng Lifeact-MS2-YFP. Gag aggregated with linear F-actin bundles at or near the cell surface even at early time points (T=0).

### Targeting *gag* mRNAs to non-PM membranes or the actin cytoskeleton alters sites of virus particle assembly

Because the MS2-YFP proteins were proxies for MSL-bearing RNAs, it was essential to confirm the subcellular localization of native *gag-pol* mRNAs using single molecule fluorescence *in situ* hybridization (smFISH) in conjunction with super-resolution structured illumination microscopy (SIM). In these 3-color experiments, we transfected HeLa cells with Rev-dependent GagFP-MSL-RRE and MS2-YFP fusion proteins either with or without Rev, fixed cells at ~30 hours post-transfection, and performed smFISH using a *gag/gag-pol*-specific DNA probe set (Fig 7A). HeLa cells transfected with MS2-YFP-NLS and GagFP-MSL-RRE in the absence of Rev exhibited marked co-localization between MS2-YFP-NLS and the gRNA FISH signal in the nucleus, with no apparent GagFP expression, consistent with robust transcription but the inability of these intron-retaining mRNAs to escape the nucleus through the CRM1 export pathway (Fig 7A, NLS no Rev condition, and S3 Video). When Rev was co-expressed, both the MS2-YFP-NLS and gRNA FISH signals shifted to the cytoplasm, consistent with nuclear export, and were now readily detected in a diffuse distribution throughout the cytoplasm as well as co-localizing with GagFP at PM-adjacent punctae, (Fig 7A, NLS+Rev condition, and S4 Video). These control experiments confirmed that the MS2-YFP signals tracked in Figs 1–6 were indeed representative of actual gRNA trafficking.

**Fig 7.**
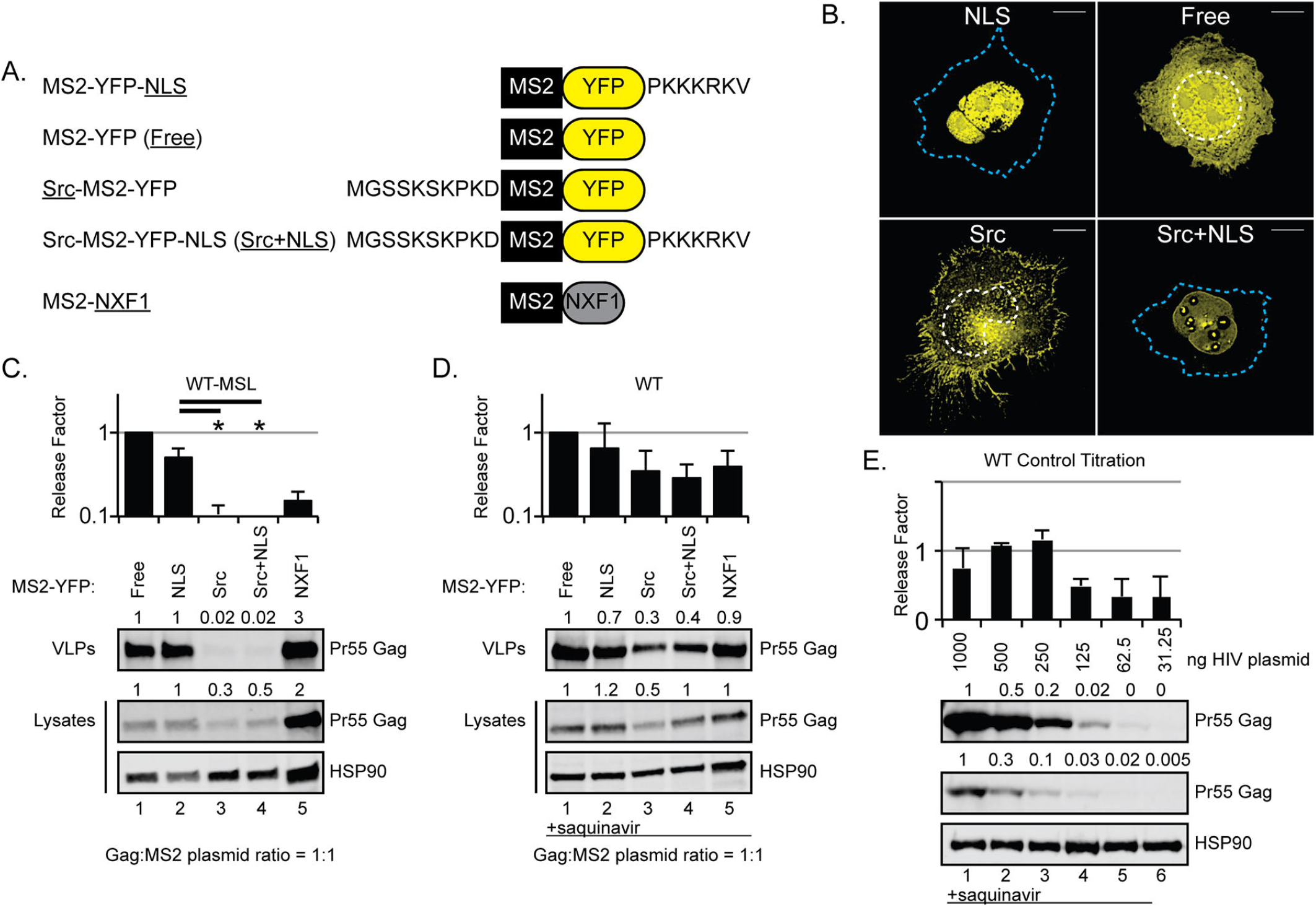
gRNA-directed effects on Gag trafficking and subcellular sites of virus particle assembly. (A) HeLa cells were transfected with 800ng GagFP-MSL-RRE, 100ng MS2-YFP targeting constructs as indicated, and 100ng Rev or empty vector as indicated. Cells were fixed at ~30 hours post-transfection, subjected to FISH, and multi-Z stack images acquired by structured-illumination microscopy (SIM) using a 100x (NA 1.49) TIRF oil objective. Single Z-plane images are shown with scale bars representing 5 microns. White arrows indicate points of interest highlighting colocalization. Nucleus = “nuc”, cytoplasm = “cyto”. (B) HEK293T cells were transfected with 1800ng WT-MSL and 200ng MS2-YFP targeting constructs as indicated for EM. Red arrows indicate representative particle events for budding (NLS), intracellular (Src), and incomplete (Lifeact). (C) Quantification of particles with budding phenotypes observed by thin-section EM. Bar graph shows percent of assembly events exhibiting intracellular, incomplete, or budding phenotype. Errors bars represent standard deviation from the mean of 10 cells imaged. At least 100 budding events were quantified per condition.

As expected, Src-MS2-YFP also co-localized with the gRNA smFISH signal. However, for this condition the Src modification triggered a massive relocalization of gRNAs from a diffuse cytoplasmic distribution to perinuclear membranes including, apparently, the nuclear envelope itself (Fig 7A, Src+Rev condition, and S5 Video). Consistent with the time lapse imaging presented in Fig 4B, we observed notable accumulations of GagFP at or near the perinuclear sites (Fig 7A, compare blue panels for NLS+Rev and Src+Rev conditions). Moreover, thin section electron microscopy on HEK293T cells transfected with WT-MSL and MS2-YFP targeting constructs confirmed a high frequency of intracellular VLP assembly events for the Src-MS2-YFP condition relative to the MS2-YFP-NLS control (50% of particles detected in association with intracellular vesicles, Fig 7B and quantification in 7C). Thus, gRNA-directed Gag trafficking to the “wrong” cellular membranes was likely to explain the bulk of the virus particle release defect observed for the Src-MS2-YFP RNA tether. Interestingly, Lifeact-MS2-YFP clearly did not abolish Gag’s trafficking to the PM (Figs 7A and S6 Video). However, we observed a large number of partially budding structures at the cell surface for this condition (defined as electron dense shells less than 75% complete) (Figs 7B and 7C). Thus, Gag tethered to actin filaments through its mRNA may be sufficient to confer an assembly defect. Taken together, these high resolution imaging strategies provided further confirmation that redirecting *gag* mRNAs to aberrant sites in the cytoplasm can markedly affect Gag subcellular distribution and reduce virus particle output.

### The *gag-pol* mRNA’s capacity to influence Gag trafficking maps to both Psi and the RRE

That *gag* mRNA trafficking influences the subcellular distribution of its protein product is consistent with a model wherein localized translation allows gRNAs to help compartmentalize Gag to the bud site. gRNAs encode two well-characterized *cis*-acting trafficking elements, Psi encoded within the 5’UTR is bound by Gag to facilitate gRNA encapsidation [22,33] and the RRE that is essential for the nuclear export of gRNAs and other intron-retaining mRNAs [57,58] (e.g., see Fig 8A). We compared the effects of MS2 targeting proteins at relatively low MS2:gRNA plasmid ratios (1:17) on WT *gag* mRNAs, a version mutated to no longer encode Psi (dPsi.GagFP-MSL-RRE), and a version that retained Psi but was rendered Rev-independent by replacing the RRE with four copies of the constitutive transport element (CTE) derived from Mason-Pfizer monkey virus that is well known to direct mRNA nuclear export to the NXF1/NXT1 pathway (depicted in Fig 8A) [77,101,102]. To our surprise, neither Src-MS2-YFP nor Lifeact-MS2-YFP had a negative effect on Gag derived from the dPsi mutant transcript (compare Fig 8B to 8C), and exerted only mild effects for the CTE condition (Fig 8D).

**Fig 8.**
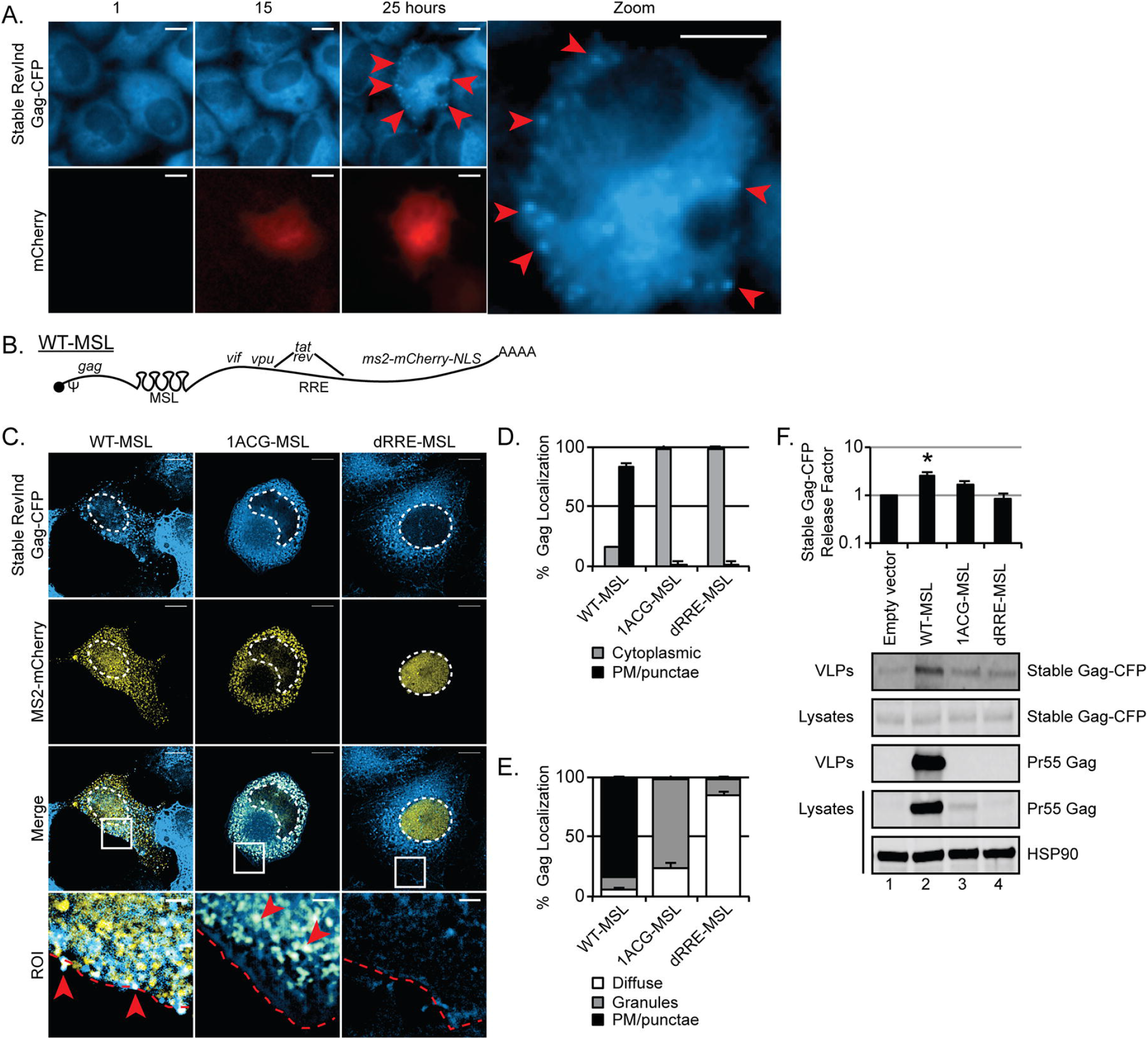
The HIV-1 *gag-pol* mRNA’s capacity to regulate Gag trafficking maps to both Psi and the RRE. (A) Cartoon depiction of constructs used in these studies. 4xCTE = four copies of the constitutive transport element from Mason-Pfizer monkey virus. (B-E) HEK293T cells were transfected with 100ng MS2-YFP targeting constructs as indicated and 1700ng of GagFP-MSL-RRE and 200ng pRev (B), 1700ng dPsi.GagFP-MSL-RRE and 200ng pRev (C), 1700ng GagFP-MSL-CTE and 200ng empty vector (D), or 1700ng RevInd GagFP-MSL and 200ng empty vector (E). VLPs and cell lysates were immunoblotted for Gag and HSP90. Bar graphs show release factor relative to MS2-YFP-NLS. Error bars represent standard deviation from the mean of three independent experiments. The asterisks (*) indicate GagFP release factor is significantly different for comparisons indicated by black bars (two-tailed Student’s t-test, p=0.018 Src & 0.047 Lifeact). (F) Cartoon depiction of RevInd GagFP constructs used in (G). (G) HEK293T cells were transfected with 100ng MS2-YFP targeting constructs as indicated, 1700ng of RevInd GagFP constructs as indicated, and either 200ng empty vector or pRev as indicated and immunoblotted for Gag and HSP90.

We also tested a dual Psi- and RRE-minus condition, using a transcript encoding codon-optimized, RevInd GagFP (as for Fig 1A) but augmented to carry the 24xMSL cassette in the 3’UTR (depicted in Fig 8A). Interestingly, both Src-MS2-YFP and Lifeact-MS2-YFP actually enhanced VLP production for Gag derived from these transcripts (Fig 8E and 8G, lanes 1–6). Genetic manipulation of this construct allowed us to test the sufficiency of either RNA element (Psi and/or RRE) to affect Gag trafficking. Gag derived from RevInd GagFP-MSL constructs modified to bear the 5’UTR region encompassing Psi (RevInd Psi-GagFP-MSL, shown in Fig 8F) became highly sensitive to Src-MS2-YFP and LifeAct-MS2-YFP expression both in terms of cytopolasmic abundance of Gag and net VLP production (Fig 8G, compare lanes 7-9 to 10-12). Remarkably, Gag derived from RevInd transcripts modified to bear the RRE (RevInd GagFP-MSL-RRE, shown in Fig 8F) also became sensitive to the Src-MS2-YFP targeting protein, but only when expressed in the presence of Rev (Fig 8G, compare lanes 13-15 to 16-18). Therefore, either Psi or the RRE (in the presence of Rev) structures were sufficient for Gag synthesis and/or assembly to be affected by targeting mRNAs to non-PM locations.

## DISCUSSION

For retroviruses, gRNA nucleocytoplasmic transport is a tightly regulated process ensuring robust late gene expression and efficient genome encapsidation during virion assembly. Herein we provide, to our knowledge, the first direct evidence that gRNA subcellular distribution represents a core determinant of the HIV-1 virion assembly pathway. We initially hypothesized that increasing the net abundance of PM-proximal gRNAs would stimulate virus particle assembly, according to the assumption that gRNAs encode one or more signals relevant to the nucleation of the assembly event at the PM [13,32,41]. However, increasing levels of “Gag-minus” 1ACG-gRNAs in *trans* had little to no effect on assembly either at high (Fig 1) or low (Fig 2) levels of GagFP. In fact, 1ACG-gRNAs arrested GagFP in cytoplasmic granules in our stable “low” GagFP cell line, suggesting that a suboptimal Gag-gRNA stoichiometry is detrimental to the formation and transit of gRNP trafficking complexes. Efforts to bias gRNP trafficking to the PM using our MS2-based RNA tethering strategy were also not beneficial to assembly, but instead inhibited virus particle production for Gag derived from gRNAs as well as *Psi*- or Rev/RRE-bearing *gag* mRNAs (Figs 3–8). Time resolved imaging confirmed striking changes to Gag subcellular distribution in living cells (Figs 5B and 6C), and single-molecule FISH coupled to super-resolution microscopy in fixed cells clearly demonstrated that both the Src-MS2-YFP and Lifeact-MS2-YFP proteins markedly altered the distribution of MSL-gRNAs and Gag away from their native, “diffuse” cytoplasmic pattern to accumulate preferentially at subcellular membranes or in association with F-actin, respectively (Fig 7A and S5 Video and S6 Video).

Although artificial, the MS2 tethering strategy provides for comparative measurements of mRNA trafficking, translation, and Gag function in single cells, thus providing useful insights relevant to the native assembly pathway. That mRNA-linked effects on Gag trafficking and assembly were only observed for gRNAs or *gag-pol* mRNAs competent for Gag synthesis (Fig 4D) bearing either Psi or the RRE *cis*-acting structural elements (Fig 8) suggests a *cis*-biased model for assembly wherein coordination of gRNA nuclear export, gRNA stability in the cytoplasm, localized translation, and compartmentalized Gag-gRNA interactions regulate Gag’s assembly efficiency at the PM (working model presented in Fig 9). Consistent with a model for localized translation, we have detected ribosomal proteins (*e.g.*, RPL9-YFP) at or near Gag/gRNA complexes both at the PM and at “re-targeted” vesicular and F-actin locations (Fig 9B). However, additional work is necessary to directly assess if these sites are truly active for Gag translation and, if so, for how long.

**Fig 9.**
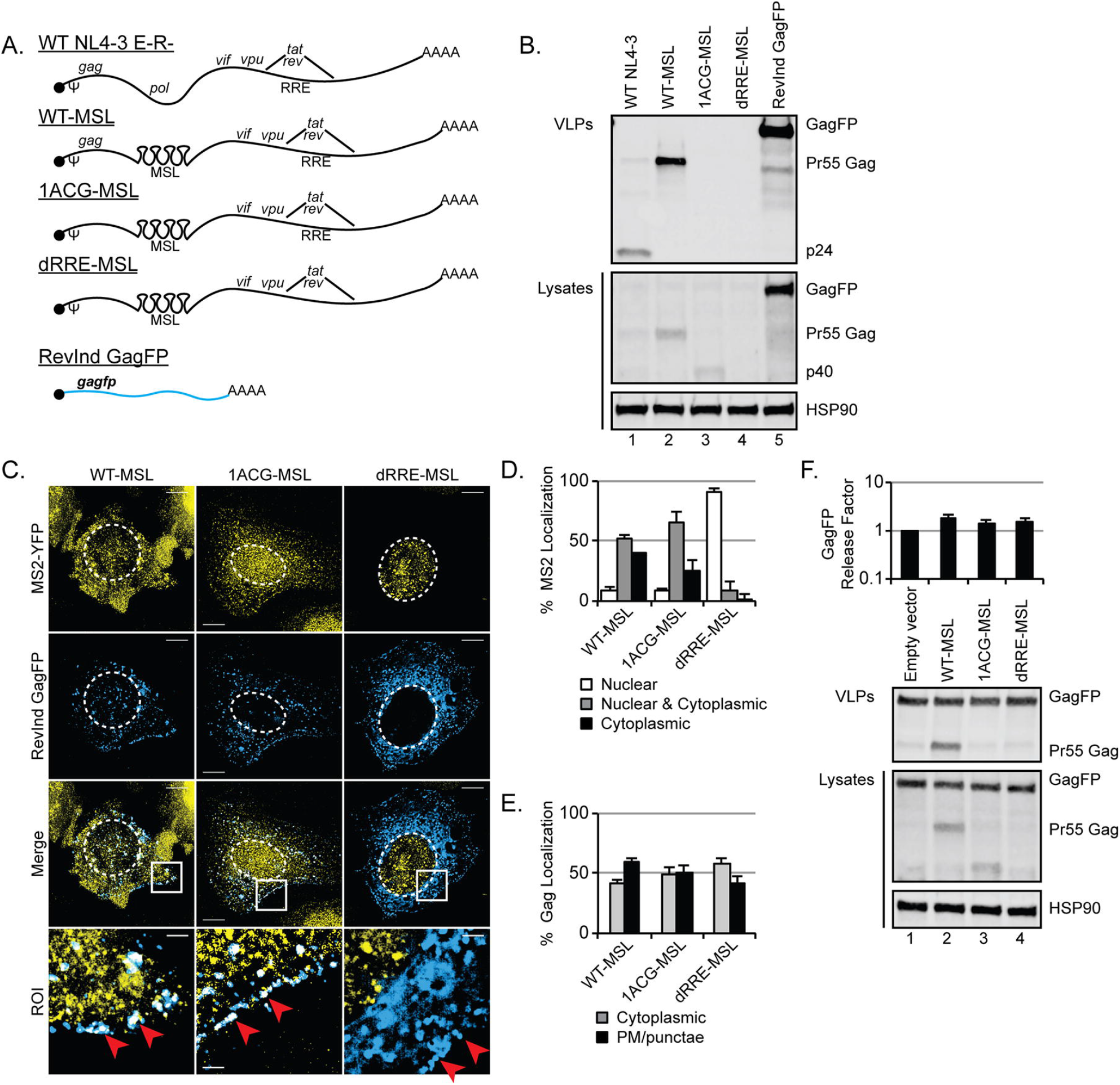
Right place, right time model for HIV-1 gRNA trafficking, Gag translation, and virion assembly. (A) HIV-1 gRNAs (red line) are exported from the nucleus via the RRE/Rev/CRM1 pathway. Once in the cytoplasm, the HIV-1 gRNA are translated (purple ribosomes) to yield some amount of the Gag polyprotein. We suggest that these HIV-1 gRNAs (likely bound by a small amount of Gag, possibly a Gag dimer) freely diffuse toward the PM where gRNA dimerization, further Gag translation, high-order Gag multimerization, increased Gag-gRNA interactions, and virion assembly occur. If these gRNAs are redirected to different subcellular locales (“Bad Neighborhoods”) such as intracellular vesicles (tan circles) or actin filaments (black chalk lines), Gag trafficking is similarly redirected and virion assembly is subsequently inhibited. In our system, using Src-MS2-YFP and Lifeact-MS2-YFP we have observed this redirection and inhibition that occurs via *cis* interactions between Gag and gRNA and is RRE/Rev-dependent. It is possible that gRNAs exported through the CRM1 pathway are marked, restructured, or coated by some as of yet unknown “missing link” indicated here by pink factor X. (B) Consistent with our model, ribosomal proteins (RPL9-YFP, yellow) were observed at PM-adjacent punctae colocalizing with Gag-CFP (cyan), and MS2-tagged gRNA (MS2-mCherry-NLS, magenta) in transfected HeLa cells. RPL9-YFP also colocalized with Gag-CFP in the presence of Src-MS2-mCherry and Lifeact-MS2-mCherry. Scale bars represent 10 µm in full images and 2 µm in regions of interest (ROI). White boxes outline the ROIs. White arrows indicate sites where MS2-mCherry, RPL9-YFP and Gag-CFP have accumulated. White pixels occur where three colors overlap.

Compartmentalization of Gag synthesis and gRNA binding in space and time may explain why gRNA encapsidation is so highly efficient despite Gag’s ready capacity to bind to non-viral RNAs and assemble non-infectious particles even in the absence of gRNA binding. At early time points, we speculate that Gag and gRNAs are maintained in the cytoplasm at low abundance in order to avoid nucleating the formation of cytoplasmic granules that may represent kinetically trapped (“dead-end”) complexes (Fig 2D and Movie 3). Low-order Gag-gRNA interactions may control how much Gag is translated [103], and diffusion in the cytoplasmic fluid will ultimately allow assembly intermediates to achieve close proximity to the PM [36,37,104]. That mistargeted gRNAs are capable of “dragging” Gag to (and/or translate Gag at) aberrant locales and, in some instances, perturb the efficiency of budding (note partial capsids for the Lifeact condition shown in Fig 7B) suggests a strong physical interaction. In this context, Psi’s role while bound by Gag likely explains why this element is both necessary and sufficient to allow for MS2-mediated perturbation of Gag distribution (Fig 8). That the RRE (with Rev) has a similar but less potent effect (Fig 8G) is more confounding. However, Gag was recently shown by Bieniasz and colleagues to bind the RRE with specificity [53] and Rev and/or the RRE have previously been shown to play roles in gRNA encapsidation [105,56,55,106].

We emphasize that under native conditions, it is almost certain that Gag plays the dominant role in defining the preferred site of assembly by tethering gRNAs to the PM through the activity of its N-terminal Matrix domain [13,41]. However, our model does not rule out potential contributions from one or more cellular RNA binding proteins that, like the MS2-targeting proteins, modulate transient Gag/gRNA interactions with membranes or other cellular machineries (Fig 9, factor “X”). The coating/coding of mRNAs with cellular factors influences their size, hydrophobicity, fluid phase, localization, and utilization [107,108], and a plethora of RNA binding proteins (ABCE1, DDX3, hnRNP isoforms, RHA, Staufen, SR proteins, among others) have already been implicated in the formation and maintenance of the HIV-1 Gag/gRNA trafficking granules [109,110,34,111,39,112,35,77,80,113–117]. We also note that unique spatiotemporal features of the Rev-regulated nuclear export pathway (*e.g.*, punctuated, rapid increases to free Gag/gRNP abundance in the cytoplasm, or diffusion in itself; Fig 4C) should influence the efficiency of the assembly pathway [82]. In this context, deficiencies to one or more co-factors tied to Rev-dependent trafficking dynamics or downstream cytoplasmic events may underpin previously observed changes to Gag assembly competency when its message is rendered Rev/RRE-independent [35,77,78,80,81].

In general, large vRNP complexes trafficking through the cytoplasm prior to utilization draw parallels with cellular mRNA molecules that are translated locally [118–120]. Perturbations to the ability of gRNAs to freely diffuse through a dense cytoplasmic fluid may have negative consequences for localized translation and subsequent virion assembly. To date, there are no FDA-approved antiviral approaches that perturb HIV-1 gene expression or the stages upstream of immature virus particle assembly, although several strategies have been pursued including disruption of Tat or Rev function, virus-specific miRNAs, and trans-dominant proteins [reviewed in [121]]. Indeed, the regulation RNP complex formation is of increasing interest in other diseases where malformation of aberrant RNA/protein aggregates or disruption of normal fluid phase dynamics are evident [4,122,107,123,124]. Our results suggest that strategies to successfully disrupt viral mRNA subcellular distribution, gRNP complex formation, or fluid phase transitions using small molecule inhibitors or alternative strategies (*e.g.*, provision of *trans*-acting synthetic “restriction” factors via gene therapy) merit further exploration.

## MATERIALS AND METHODS

### Cell culture, plasmids, and stable cell lines

Human HeLa and HEK293T cell lines (obtained from the ATCC) were cultured in DMEM (Sigma-Aldrich, Madison, WI, USA) supplemented with 10% fetal bovine serum, 1% L-glutamine, and 1% penicillin-streptomycin. Full-length parental WT HIV-1 proviral plasmids were derived from the pNL4-3 molecular clone [125] bearing inactivating mutations in *env*, *vpr*, and expressing a Firefly Luciferase reporter from the *nef* reading frame (E-R-/Luc) [126]. 24 copies of the MS2 bacteriophage RNA stem loop (MSL, a kind gift of Robert Singer, Albert Einstein University, New York, NY) were engineered into the full-length pNL4-3 derived constructs as previously described [127] thereby generating pNL4-3/E-R-Luc-24xMSL (WT-MSL). Replacing Firefly Luciferase reporter E-R-/Luc with mCherry in the nef reading frame using NotI and XhoI cut sites generated HIV-1/mCherry virus. Subgenomic GagFP-MSL HIV-1 expression plasmids encoded Gag fused to mTagBFP2 [128], ECFP, or mCherry upstream of the MSL cassette and inserted into surrogate, subgenomic HIV-1 gRNA plasmid Gag-Pol-Vif-RRE or Gag-Pol-Vif-4xCTE [113,82]. Rev-independent (RevInd) Gag-fluorescent protein (FP) plasmids were derived from a plasmid encoding partially codon-optimized Gag-GFP (a gift of Marilyn Resh, Memorial Sloan Ketterlng Cancer Center, New York, NY, USA) [85,86]. The FP reading frame was fused in frame to RevInd Gag cDNAs using overlapping PCR and inserted into pcDNA3.1 using *Nhe*I and *Xho*I cut sites. In all instances, mutants of full-length HIV-1 and RevInd Gag plasmids were generated using overlapping PCR. pRevInd-GagFP-MSL was generated by inserting the 24xMSL cassette into pRevInd-GagFP using *BsrGI* cut sites, Psi-RevInd-MSL GagFP by inserting HIV-1_NL4-3_ 5’UTR nts 1-336 into *Nhe*I and *Sac*II sites in pRevInd-GagFP-MSL, and pRevInd-GagFP-MSL-RRE by transferring the RevInd-GagFP-MSL sequence into a pcDNA-RRE backbone plasmid using *Sac*I and *EcoR*I sites. mTagBFP2 was a gift from Michael Davidson (Addgene plasmid # 55302). pRev has been described [77]. MS2-YFP targeting constructs were generated by amplifying cDNAs from pMS2-YFP (also a gift of Rob Singer, Albert Einstein University, New York, NY, USA) [129] using overlapping PCR prior to subcloning into a pcDNA3.1 backbone using *Hind*III and *Xho*I cut sites. MS2-mCherry-NLS was generated by overlapping PCR to replace YFP with the mCherry reading frame and subcloned into the *nef* position of the full-length pNL4-3/E-R-Luc-24xMSL HIV constructs using *Not*I and *Xho*I cut sites. MS2-YFP targeting constructs included an amino-terminal membrane targeting signal derived from the Src kinase (MGSSKSKPKD) [87], amino-terminal Lifeact actin-targeting domain (MGVADLIKKFESISKEE) [100], and/or a carboxy-terminal nuclear localization signal (NLS; PKKKRKV) derived from the SV40 Large T antigen [130]. pRPL9-YFP was subcloned from human cDNA using overlapping PCR. HeLa.MS2-YFP, HeLa.Gag-CFP, and HEK293T.Gag-CFP stable cell lines were generated as previously described [131–133,82]. Briefly, MS2-YFP or Gag-CFP reading frames were subcloned into a MIGR1-derived retroviral vector (pCMS28) upstream of sequence encoding an internal ribosomal entry site (IRES) regulating a second reading frame encoding Puromycin-N-acetyltransferase [131]. High performance clones were selected by limiting dilution in 2µg/mL puromycin.

### Retroviral assembly assays

Cells at 30-40% confluency were transfected with 2µg DNA in six well dishes using polyethylenimine (PEI; #23966, Polysciences Inc, Warrington, PA, USA). pcDNA3.1 or pBlueScript were used as empty vector controls. Culture media were replaced at 24 hours post-transfection and cell lysates and supernatants were harvested for immunoblot analysis at 48 hours as previously described [80]. Briefly, 1mL of harvested culture supernatant was filtered, underlaid with 20% sucrose (w/v) in PBS, subjected to centrifugation at >21,000g for two hours at 4ºC, and viral pellets were resuspended in 35µL dissociation buffer (62.5 mM Tris-HCl, pH 6.8, 10% glycerol, 2% sodium dodecyl sulfate [SDS], 5% β-mercaptoethanol). Cells were harvested in 500µL radioimmunoprecipitation assay (RIPA) buffer (10 mM Tris-HCl, pH 7.5, 150 mM NaCl, 1 mM EDTA, 0.1% SDS, 1% Triton X-100, 1% sodium deoxycholate), lysed by passage through a 26G needle, subjected to centrifugation at 1,500g for 20 minutes at 4ºC, and combined 1:1 with 2X dissociation buffer. Proteins were resolved by sodium dodecyl sulfate-polyacrylamide gel electrophoresis (SDS-PAGE) and transferred to nitrocellulose membranes. Gag was detected using a mouse monoclonal antibody recognizing HIV-1 capsid/p24 (183-H12-5C; 1:1000 dilution) from Dr. Bruce Chesebro and obtained from the NIH AIDS Research and Reference Reagent Program (Bethesda, MD, USA) [134] and anti-mouse secondary antibodies conjugated to an infrared fluorophore (IRDye680LT, 1:10000 dilution, Li-Cor Biosciences, Lincoln, NE, USA) for quantitative immunoblotting. As a loading control, heat shock protein 90A/B (HSP90) was detected using a rabbit polyclonal antibody (H-114, 1:2500 dilution, Santa Cruz Biotechnology, Santa Cruz, CA, USA) and anti-rabbit secondary antibodies conjugated to an infrared fluorophore (IRDye800CW, 1:7500 dilution, Li-Cor Biosciences). Where indicated, the protease inhibitor saquinavir (NIH AIDS Research and Reference Reagent program, Bethesda, MD) was added at 24 hours post-transfection. Typically, retroviral assembly assays were performed with transfections and harvesting occurring in one week while processing and immunoblotting occurred in the following week. To ensure reproducibility, most results were obtained from three biological replicates as defined as cells plated in six well dishes transfected on separate days (*i.e.* replicate 1 was transfected on a separate day from replicate 2).

### Microscopy and fluorescence in situ hybridization (FISH)

Cells were plated in 24-well glass-bottom dishes (Mattek Corporation, Ashland, MA, USA) or 8-well microslides (IBIDI, Madison, WI, USA) and transfected using PEI. Transfection mixes contained 1µg (24-well) or 333ng (IBIDI) plasmid DNA, respectively. Deconvolution fixed-cell imaging experiments were performed on a Nikon Ti-Eclipse inverted wide-field microscope (Nikon Corporation, Melville, NY, USA) using a 100x Plan Apo oil objective lens (numerical aperture NA 1.45). These cells were fixed 24-32 hours post-transfection in 4% paraformaldehyde in PBS. Live cell imaging experiments were also performed on a Nikon Ti-Eclipse inverted wide-field microscope using a 20x Plan Apo objective lens (NA 0.75) with images acquired typically every 60 minutes over a time course of 16-36 hours. Images were acquired using an ORCA-Flash4.0 CMOS camera (Hamamatsu Photonics, Skokie, IL, USA) and using the following excitation/emission filter sets (nanometer ranges): 430/470 (CFP), 510/535 (YFP), 585/610 (mCherry).

For fixed cell experiments using smFISH to visualize HIV-1 gRNA, cells were plated and transfected as above. At ~30 hours post-transfection, cells were washed, fixed in 4% formaldehyde, and permeabilized in 70% ethanol for at least four hours at 4ºC. Custom Stellaris FISH probes were designed to recognize NL4-3 HIV-1 *gag-pol* reading frame nucleotides 386-4614 by utilizing Stellaris RNA FISH Probe Designer (Biosearch Technologies, Inc., Petaluma, CA, USA) available online at www.biosearchtech.com/stellarisdesigner (version 4.1). The samples were hybridized with the Gag/GagPol Stellaris RNA Fish Probe set (48 probes) labeled with CAL Fluor Red 610 dye (Biosearch Technologies, Inc.), following manufacturer’s instructions available online at www.biosearchtech.com/stellarisprotocols. Structured illumination microscopy (SIM) was performed on a Nikon N-SIM microscope using a 100x TIRF oil objective lens (NA 1.49). Images were acquired using an Andor iXon Ultra 897 EMCCD (Andor Technology, Belfast, United Kingdom) and Nikon NIS Elements in 3D-SIM mode using the following excitation laser wavelengths (nanometer ranges): 408 (mTagBFP2), 488 (YFP), and 561 (CAL Fluor Red 610). Widefield epifluorescent microscopy images were deconvolved using NIS Elements. All images were processed and analyzed using FIJI/ImageJ2 [135]. Results were obtained from three biological replicates as defined as cells plated in IBIDI slides or 24-well dishes transfected on separate days *(i.e.* replicate 1 was transfected on a separate day from replicate 2).

### Thin section electron microscopy

For thin section EM, HEK293T cells were cultured in six-well dishes and transfected as described above and processed as previously described [136]. At 48 hours post-transfection, cells were fixed in a solution of 2.5% glutaraldehyde, 2.0% paraformaldehyde in 0.1M sodium phosphate buffer (PBS), pH 7.4 for ~2 hours at room temperature. Samples were rinsed five times for five minutes each in 0.1M PBS. Rinsed cells were post-fixed in 1% osmium tetroxide, 1% potassium ferrocyanide in PBS for 1 hour at room temperature. Following osmium tetroxide post-fixation, the samples were rinsed in PBS, as before, and rinsed three times in distilled water for five minutes to clear phosphates and embedded using increasing concentrations (10mL A/M, 10mL B, 300µL C, 100µL D components) of Durcupan ACM resin (Fluka AG, Switzerland) at 60ºC. Cells were pelleted and sectioned using a Leica EM UC6 ultramicrotome with 100nm sections collected on 300 mesh copper thin-bar grids, and contrasted with Reynolds lead citrate and 8% uranyl acetate in 50% ethanol. Sections were observed with a Phillips CM120 transmission electron microscope, and images were collected with a MegaView III (Olympus-SIS, Lakewood, CO, USA) side-mounted digital camera. All images were processed and analyzed using FIJI/ImageJ2 [135].

## ACKNOWLEDGEMENTS

We gratefully acknowledge Randall Massey and the University of Wisconsin SMPH Electron Microscopy Facility for assistance with processing and imaging EM. We gratefully acknowledge Elle Kielar Grevstad and the University of Wisconsin Biochemistry Optical Core for assistance with structured illumination microscopy. The following reagents were obtained through the NIH AIDS Reagent Program, Division of AIDS, NIAID, NIH: HIV-1 p24 Hybridoma (183-H12-5C) (from Dr. Bruce Chesebro) and Saquinavir. We wish to thank Bill Sugden, Sam Butcher, Halena VanDeusen, Ed Evans, and Bayleigh Benner for their assistance in reading this manuscript and helpful intellectual discussions.

## SUPPLEMENTARY MATERIAL

**S1 Video. HIV-1 infection drives stably-expressed GagFP to assembly sites at the PM.** Live cell imaging of HeLa.Gag-CFP (cyan) cells infected with WT NL4-3 E-R-/mCherry reporter virus. Images were acquired once per hour and are shown here at 4 frames per second. Scale bar represents 10 microns. White arrows indicate sites where stably expressed RevInd Gag-CFP has accumulated in PM-adjacent punctae.

**S2 Video. HIV-1 “Gag-minus” 1ACG-gRNAs trigger accumulation of stably-expressed GagFP into cytoplasmic granules.** Live cell imaging of HeLa.Gag-CFP (cyan) cells transfected with 1ACG-MSL/MS2-mCherry (yellow). Images were acquired once per hour and are shown here at 4 frames per second. Scale bar represents 10 microns. White arrows indicate sites where stably expressed RevInd Gag-CFP and MS2-mCherry-tagged gRNA have accumulated in cytoplasmic granules.

**S3 Video. Three-dimensional view of MS2-YFP-NLS and smFISH tagged HIV-1 gRNA expressed in the absence of Rev.** SIM image reconstructed in 3D using FIJI/ImageJ2. Scale bar represents 5 microns. MS2-YFP (green) and gRNA FISH (red) are sequestered in the nucleus in the absence of the Rev protein. White arrow indicates edge of nucleus (nuclear envelope).

**S4 Video. Three-dimensional view of MS2-YFP-NLS, smFISH tagged HIV-1 gRNA, and HIV-1 Gag.** SIM image reconstructed in 3D using FIJI/ImageJ2. Scale bar represents 5 microns. MS2-YFP (green) and gRNA FISH (red) colocalize in the cytoplasm and at the PM with Gag (blue). White arrows indicate colocalized MS2-YFP, gRNA, and Gag punctae at PM.

**S5 Video. Three-dimensional view of Src-MS2-YFP, smFISH tagged HIV-1 gRNA, and HIV-1 Gag.** SIM image reconstructed in 3D using FIJI/ImageJ2. Scale bar represents 5 microns. MS2-YFP (green) and gRNA FISH (red) colocalize at intracellular vesicles and the nuclear periphery with Gag (blue). White arrows indicate colocalized MS2-YFP, gRNA, and Gag punctae at intracellular membranes.

**S6 Video. Three-dimensional view of Lifeact-MS2-YFP, smFISH tagged HIV-1 gRNA, and HIV-1 Gag.** SIM image reconstructed in 3D using FIJI/ImageJ2. Scale bar represents 5 microns. MS2-YFP (green) and gRNA FISH (red) colocalize along linear F-actin filaments with Gag (blue). White arrows indicate colocalized MS2-YFP, gRNA, and Gag punctae along actin filaments.

**S5 Video. Three-dimensional view of Src-MS2-YFP, smFISH tagged HIV-1 gRNA, and HIV-1 Gag.** SIM image reconstructed in 3D using FIJI/ImageJ2. Scale bar represents 5 microns. MS2-YFP (green) and gRNA FISH (red) colocalize at intracellular vesicles and the nuclear periphery with Gag (blue). White arrows indicate colocalized MS2-YFP, gRNA, and Gag punctae at intracellular membranes.

**S6 Video. Three-dimensional view of Lifeact-MS2-YFP, smFISH tagged HIV-1 gRNA, and HIV-1 Gag.** SIM image reconstructed in 3D using FIJI/ImageJ2. Scale bar represents 5 microns. MS2-YFP (green) and gRNA FISH (red) colocalize along linear F-actin filaments with Gag (blue). White arrows indicate colocalized MS2-YFP, gRNA, and Gag punctae along actin filaments.

